# Temperature sensitive SMA-causing point mutations lead to SMN instability, locomotor defects, and premature lethality in *Drosophila*

**DOI:** 10.1101/832030

**Authors:** Amanda C. Raimer, Suhana S. Singh, Maina R. Edula, Tamara Paris-Davila, Vasudha Vandadi, Ashlyn M. Spring, A. Gregory Matera

**Author notes:** Department of Biology, University of Wisconsin, River Falls, WI 54022, USA.

## Abstract

Spinal muscular atrophy (SMA) is the leading genetic cause of death in young children, arising from homozygous deletion or mutation of the *SMN1* gene. SMN protein expressed from a paralogous gene, *SMN2*, is the primary genetic modifier of SMA; small changes in overall SMN levels cause dramatic changes in disease severity. Thus, deeper insight into mechanisms that regulate SMN protein stability should lead to better therapeutic outcomes. Here, we show that SMA patient-derived missense mutations in the *Drosophila* SMN Tudor domain exhibit a pronounced temperature sensitivity that affects organismal viability, larval locomotor function, and adult longevity. These disease-related phenotypes are domain-specific and result from decreased SMN stability at elevated temperature. This system was utilized to manipulate SMN levels during various stages of *Drosophila* development. Due to a large maternal contribution of mRNA and protein, *Smn* is not expressed zygotically during embryogenesis. Interestingly, we find that only baseline levels of SMN are required during larval stages, whereas high levels of protein are required during pupation. This previously uncharacterized period of elevated SMN expression, during which the majority of adult tissues are formed and differentiated, could be an important and translationally relevant developmental stage in which to study SMN function. Altogether, these findings illustrate a novel *in vivo* role for the SMN Tudor domain in maintaining SMN homeostasis and highlight the necessity for high SMN levels at critical developmental timepoints that is conserved from *Drosophila* to humans.

## INTRODUCTION

Spinal Muscular Atrophy (SMA) is the leading genetic cause of death in infants and small children, with an incidence of ∼1:7,000 live births and a carrier frequency of ∼1:50 (Prior et al., 2010; Sugarman et al., 2012; Vill et al., 2019). This progressive neuromuscular disease is characterized by alpha-motor neuron degeneration and muscle atrophy, resulting in gradual loss of motor function. SMA symptoms present within a spectrum of disease severity. Left untreated, patients with the most severe form of the disorder are unable to stand or sit upright, and do not survive past two years of age (Crawford and Pardo, 1996; Farrar et al., 2017). In contrast, milder forms of SMA are not typically diagnosed until later in life, and these patients exhibit mild motor dysfunction, living relatively normal lifespans (Alatorre-Jiménez et al., 2015; Tiziano et al., 2013).

Despite its broad spectrum of severity, SMA is a monogenic disorder that is most commonly caused by homozygous deletion of *Survival Motor Neuron 1* (*SMN1*) and a corresponding reduction in the expression of full-length Survival Motor Neuron (SMN) protein. Animal studies have shown that complete loss of SMN protein results in death *in utero* (Schrank et al., 1997). However, the presence of a paralogous gene in humans, *SMN2*, allows for the survival of affected individuals past birth (Coovert et al., 1997). The coding region of *SMN2* is identical to that of *SMN1,* save for five nonpolymorphic nucleotide differences, one of which causes skipping of exon 7 during splicing in roughly 90% of *SMN2* pre-mRNAs (Lorson et al., 1999). Transcripts produced by this alternative splicing event are translated into a truncated version of SMN protein (SMNΔ7) and are quickly degraded by the proteasome (Gray et al., 2018; Lorson et al., 1998). The remaining fraction of full-length transcripts (∼10%) encodes full-length SMN that is identical to protein produced by *SMN1*. In humans, *SMN2* is located on chromosome 5q within a highly dynamic genomic region that is prone to both duplications and deletions (Lefebvre et al., 1995). This has led to significant *SMN2* copy number variation in the population (Butchbach, 2016; Carpten et al., 1994; Courseaux et al., 2003). Complete loss of *SMN2* has no phenotypic effect in healthy individuals; however, in SMA patients *SMN2* is the primary genetic modifier of disease severity (Feldkötter et al., 2002; Lefebvre et al., 1997; Velasco et al., 1996). Higher *SMN2* copy number produces increased levels of full-length SMN protein, which corresponds to later disease onset and milder symptoms. Although the precise molecular etiology of SMA remains unclear, overwhelming evidence shows that reduced SMN protein levels cause the disease (Ahmad et al., 2016; Briese et al., 2005; Chaytow et al., 2018; Deguise and Kothary, 2017; Li et al., 2014).

The importance of SMN protein levels is further evidenced by the fact that the mechanism of action for both FDA-approved treatments currently available for SMA, Spinraza (nusinersen) and Zolgensma (onasemnogene abeparvovec), aim to increase SMN protein levels (Sumner and Crawford, 2018). Although these treatments have dramatically improved the prognosis of SMA patients, there are still limitations to the therapies that could be addressed using combinatorial therapies (Gidaro and Servais, 2019; Ramos et al., 2019; Sumner and Crawford, 2018). For example, it remains to be seen whether these treatments will remain effective over time and into adulthood, or if the patients might develop symptoms later in life. Additionally, given SMN’s general housekeeping function in the biogenesis of spliceosomal snRNPs (Matera and Wang, 2014), long-term treatment of the CNS may reveal deficits in peripheral tissues over time. Thus, a multi-pronged approach to precisely control SMN levels and function across tissues is more likely to prevent SMA disease progression throughout a patient’s lifetime.

Although most SMA patients carry a homozygous deletion of *SMN1*, 5% of those affected are heterozygous, harboring a deletion of *SMN1* over a small indel or missense mutation (Lefebvre et al., 1995; Wirth, 2000). To better understand how *SMN1* missense mutations contribute to disease, our laboratory has developed *Drosophila* as an SMA model system. Previously, we generated an allelic series of transgenic fly lines that express SMA-causing point mutations in an otherwise *Smn* null mutant background (Praveen et al., 2012; Praveen et al., 2014). These animals express Flag-tagged wildtype or mutant SMN from the native *Smn* promoter (Fig 1A) and have been used to study SMA phenotypes at the behavioral, physiological, and molecular levels (Garcia et al., 2013; Garcia et al., 2016; Gray et al., 2018; Praveen et al., 2014; Spring et al., 2019).

**Fig. 1.**
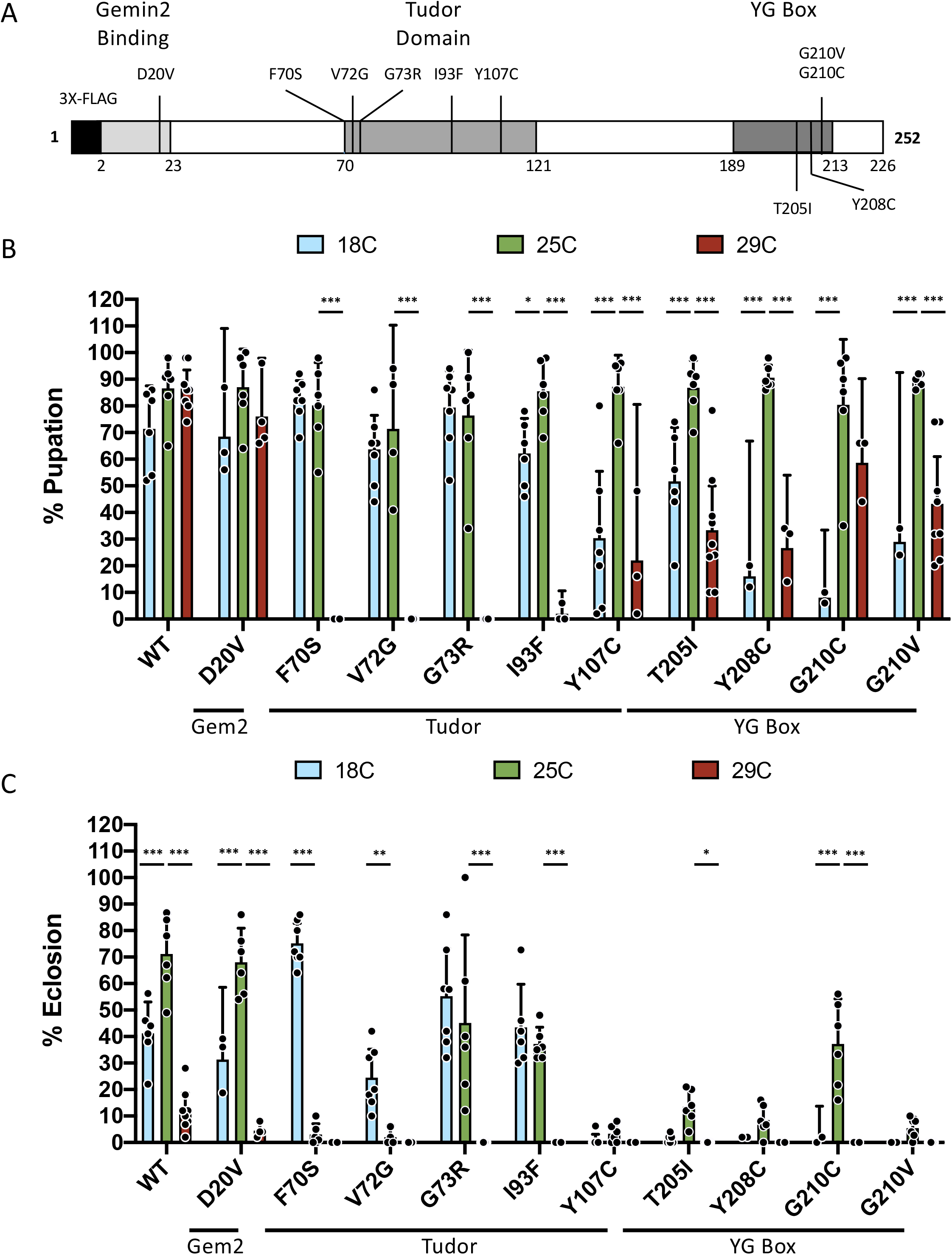
Effects of temperature limits on viability of SMN patient-derived missense mutant lines. (A) Diagram of the SMN protein showing orthologous SMA patient-derived missense mutations in the *Drosophila* SMN protein sequence. This schematic represents our *Drosophila* SMN transgene construct, which includes an N-terminal 3X-FLAG tag and a 3kB upstream sequence that includes the native *Smn* promoter. (B,C) Viability assay of the wildtype and 10 *Smn* mutant transgenic lines, measured by percentage of larvae reaching the pupal (B) and adult (C) stages when being raised at either the optimal culturing temperature, 25°C (green), or one of the extreme temperatures, 18°C (light blue) or 29°C (red). Mutants are separated by the protein functional domain that they impact (Gemin2, Tudor, YG box). The number of larvae for each genotype-temperature combination ranges from 100 to 400 larvae. Larvae were split into vials of ∼50 and the viability of each vial was treated as a separate replicate (data points). Error bars represent mean±95% c.i. Adjusted P-value was calculated using two-way ANOVA and Tukey’s multiple comparisons test. *p<0.05, **p<0.01, ***p<0.001.

SMN contains three conserved regions, including the N-terminal Gemin2 binding motif, the C-terminal YG-box self-oligomerization module and the centrally located Tudor domain. The presence of disease-causing mutations within each of these three regions demonstrates the importance of each domain to SMN function. Previous work in *Drosophila* and other models has demonstrated a functional role for the YG-box in targeting SMNΔ7 for degradation by the proteasome (Cho and Dreyfuss, 2010; Gray et al., 2018). In contrast, very little is known about the effect of the SMN Tudor domain on SMN protein levels. The canonical function of a Tudor domain is to bind to methylated arginine or lysine residues, thereby modifying the activity or function of the target protein (Pek et al., 2012). In the context of SMN, the Tudor domain binds dimethylated arginines on Sm proteins (Brahms et al., 2001; Bühler et al., 1999). This interaction assists in assembly and formation of Sm-class small nuclear ribonucleoproteins (snRNPs) (Gonsalvez et al., 2007; Gonsalvez et al., 2008; Meister et al., 2001; Pellizzoni et al., 2002). Patient data along with *in vitro* and *in silico* studies indicate that certain Tudor domain mutations affect SMN protein levels, but the mechanism underlying this phenomenon remains unclear, especially *in vivo* (Hossain and Hosen, 2019; Li, 2017; Sneha et al., 2018; Takarada et al., 2017; Tripsianes et al., 2011).

Here, we present evidence that point mutations in the SMN Tudor domain are temperature sensitive relative to the WT protein, destabilizing SMN at high temperatures. Additionally, we demonstrate that this added degree of instability reduces SMN protein levels sufficiently to impact SMA-related phenotypes such as organismal viability, larval locomotion, and adult longevity. The temperature-sensitive nature of these mutants also provides a useful experimental system in which to study how changes in SMN protein levels affect molecular and physiological processes across animal development. Collectively, the results expand our understanding of the mechanisms that not only govern SMN protein stability but also SMA etiology.

## RESULTS

### SMN Tudor domain mutants (TDMs) are temperature-sensitive

Previous work using SMA patient-derived *Smn* missense mutations, modeled in the fly, has produced robust and reproducible findings. However, in one or two instances, we noticed inconsistencies in the overall viability of a given fly line that could not be attributed to normal biological noise. For example, in Praveen et al. (2014), the *Smn^F70S^* mutation line (hereafter F70S) displayed a relatively mild phenotype, with an eclosion frequency similar to that of the *Smn^WT^* transgene (hereafter WT). In contrast, Spring et al. (2019) reported a rather severe viability defect for this same F70S line. The husbandry conditions used in each study were slightly different; the experiments in the earlier work were performed at room temperature (∼22°C) whereas in the subsequent experiments, animals were kept at a constant 25°C. In addition, we qualitatively observed that certain *Smn* mutant lines displayed a dramatic decrease in viability at 29°C compared to culturing at 25°C. These two observations led us to characterize the mechanism underlying this sensitivity, as determinants of SMN function could be of translational value to SMA patients.

Therefore, we examined the effects of temperature on the viability of eleven different SMA patient-derived mutant lines (Fig. 1A) at two temperature extremes of *Drosophila* husbandry, 18°C and 29°C, as well as at the standard condition of 25°C. Viability of each transgenic line was calculated as the fraction of animals that either pupated (% pupation, larval-to-pupal transition) or eclosed (% eclosion, pupal-to-adult transition) (Fig. 1B,C). A majority of our SMA models displayed the expected trend, with the highest viability observed at 25°C and reduced viability at the extremes. In contrast, all but one of the SMN Tudor domain mutant lines (TDMs) displayed an inverse correlation between viability and temperature. That is, lower temperatures increased viability in TDMs relative to 25°C, whereas higher temperatures decreased viability. The decrease in pupation frequency for the TDMs was dramatic and statistically different from the moderate decrease observed for mutations in other regions of SMN (Fig. 1B). Similarly, only the TDM lines displayed increased eclosion frequencies at 18°C compared to 25°C (Fig. 1C). Collectively, these data indicate that the F70S, V72G, G73R, and I93F mutations in the SMN Tudor domain are temperature-sensitive (ts) alleles.

### SMN Tudor domain mutants display SMA-related phenotypes in response to small changes in temperature

We next examined the effect of minor changes in temperature by raising animals at 22°C and 27°C. The SMN WT and T205I YG box domain mutant lines were used as controls. Although the changes in temperature were relatively small (±2-3°C), the effects on the TDMs viability were still substantial (Fig 2A,B). The WT and T205I control lines still display some variability in pupation and eclosion percentages. In contrast, although a percentage of all the TDMs do manage to pupate to at all temperatures, they again show a drastic decrease in viability that inversely correlates with temperature.

**Fig. 2.**
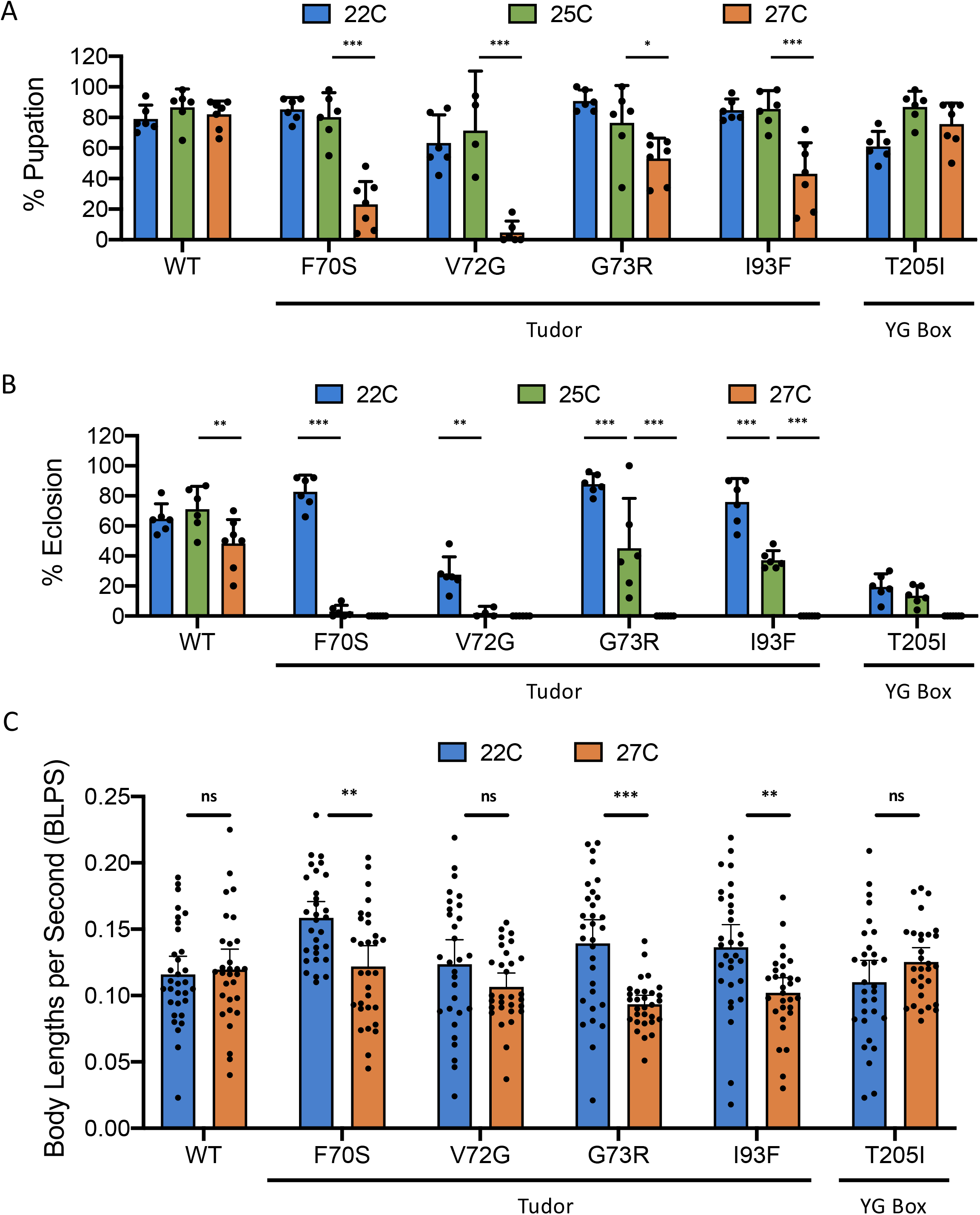
Effects of small temperature changes on SMN Tudor mutant viability and locomotor function. (A,B) Viability assay of flies expressing a wildtype *Smn* transgene, one of four Tudor domain mutations (F70S, V72G, G73R, and I93F), or a YG box domain mutation (T205I). Viability is measured as the percent of larvae that reach the pupal (A) and adult (B) stages while being raised at either the optimal culturing temperature, 25°C (green), or a slightly colder (22°C, blue) or slightly warmer (27°C, orange) temperature. The number of larvae for each experimental group ranges from 150 to 400 larvae. Larvae were split into vials of ∼50 and the viability of each vial was treated as a separate replicate. The 25°C viability data are the same data from Fig. 1C,D and are included for ease of comparison. Error bars represent mean±95% c.i. Adjusted P-values calculated using Tukey’s multiple comparisons test. *p<0.05, **p<0.01, ***p<0.001. (C) Locomotion assays of wandering 3^rd^-instar larvae of the same genotypes as (A) and (B) raised at 22°C (blue) or 27°C (orange). 30-35 larvae were assayed per condition. Locomotion was measured in body lengths per second (BLPS), which measures the larva’s speed in relationship to its body size. Error bars represent mean±95% c.i. P-values for calculated using student’s t-test. ns: not significant (p >0.05), ** p<0.01, *** p<0.001.

Having established the temperature-sensitive nature of the TDMs, we next sought to examine more SMA-related phenotypes such as muscle function indirectly through locomotor ability. Although the most extreme temperatures, 18°C and 29°C, showed the most drastic changes in viability, certain limitations make these temperatures suboptimal for analyzing *Drosophila* larval locomotion. Proper developmental staging of larvae in many assays is critical so that changes observed between wildtype and TDMs are solely attributed to the mutation and not different developmental stages (Garcia et al., 2013). Because TDMs raised at 25°C die sometime during pupation, the wandering 3^rd^-instar larval stage is one of the most developmentally significant and technically tractable stages to use when performing larval assays. However, most TDMs do not reach the wandering 3^rd^-instar at 29°C, and viability of the WT transgenic line is markedly affected at both 18°C and 29°C. In contrast, culturing at 22°C and 27°C allows for selection at wandering 3^rd^-instar stage as well as stable wildtype viability.

Larval locomotion assays were carried out on animals raised at 22°C and 27°C, as described previously (Spring et al., 2019). Crawling speed was expressed in terms of body lengths per second (BLPS), which provides a comparable measure of larval speed regardless of body size (Fig. 2C). Larvae expressing either WT or T205I SMN (a YG box mutant) display no significant changes in locomotion when raised at 22°C versus 27°C. In contrast, TDM animals raised at 22°C showed significantly improved locomotor function compared to their counterparts raised at 27°C (Fig 2C). These results highlight the fact that neuromuscular phenotypes are exacerbated specifically in the TDMs raised at elevated temperatures.

### Reduced SMN levels caused by instability underlie TDM sensitivity to temperature

Small perturbations in SMN protein levels are known to cause stark changes in patient disease severity (Lefebvre et al., 1997). We therefore evaluated SMN protein levels as a potential cause of the temperature sensitivity observed in the TDMs. SMN point mutation lines bearing a single copy of the *Smn* transgene were raised at 22°C, 25°C, or 27°C, and the SMN protein levels of wandering 3^rd^-instar larvae were measured by western blotting. Band intensities were then normalized against the total amount of protein in each sample (Fig. S1). Representative blots showing SMN protein levels in the WT and mutant larvae raised at the three different temperatures are shown in Fig. 3A. As shown, SMN levels in WT and T205I mutant animals remain relatively constant regardless of temperature. In contrast, SMN levels are markedly decreased in TDMs raised at higher temperatures. Quantification of multiple biological replicates (Fig. 3B) confirms that the trend of decreasing SMN protein levels holds for nearly all of the TDMs. Note that the very low levels of SMN in the V72G mutants raised at 25° and 27°C are close to the detection limit, so the trend is less visible in that line. Collectively, these data indicate that temperature sensitivity of TDMs is likely driven by a reduction in SMN protein levels.

**Fig. 3.**
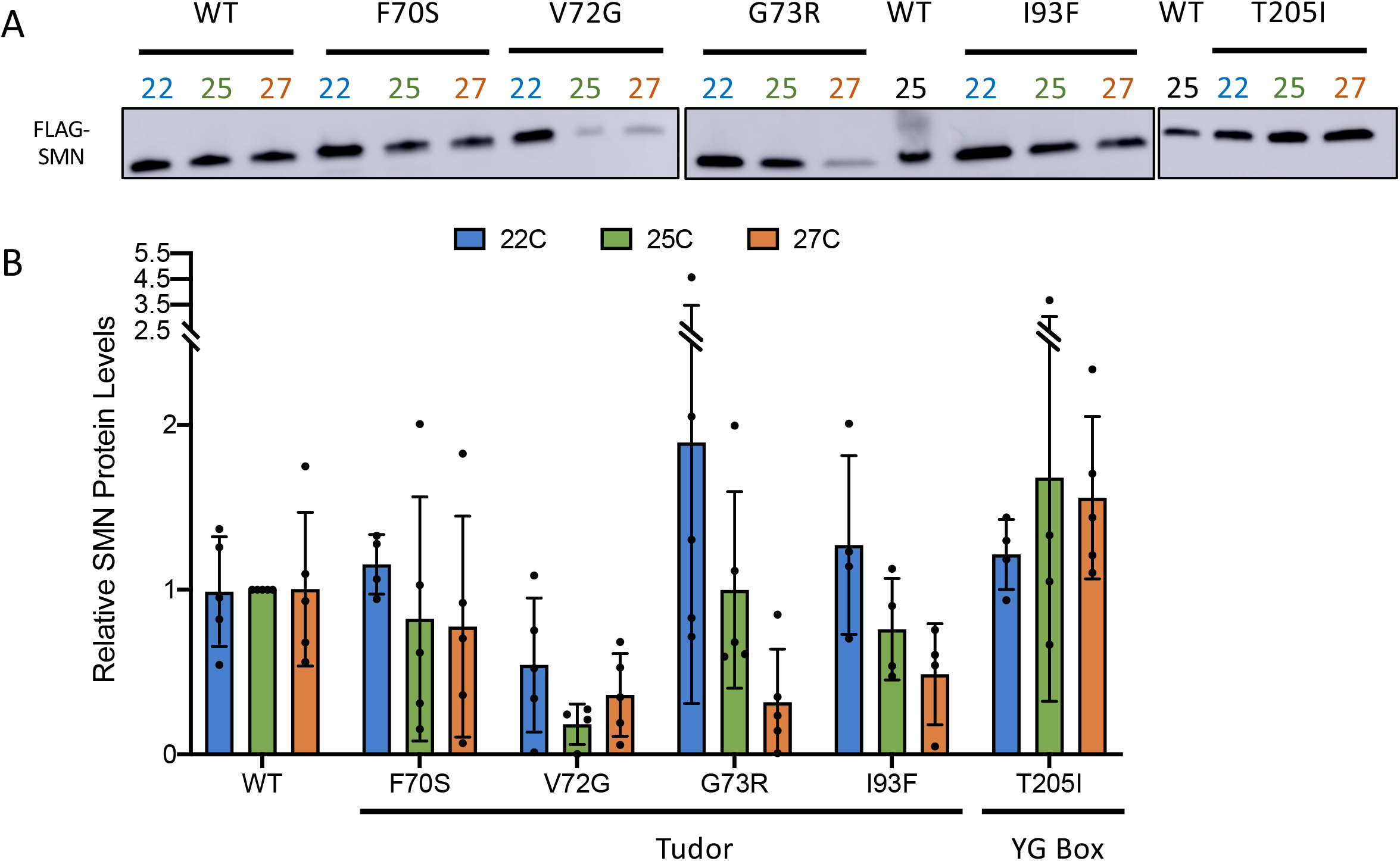
Effects of small temperature changes on SMN protein levels in Tudor domain mutants. (A) Representative Western blots of wandering 3^rd^-instar larvae from wildtype, Tudor domain mutants, and YG box domain mutant (T205I) raised at 22°C (blue), 25°C (green) or 27°C (orange), 8-12 larvae per sample. Protein was visualized using an HRP-conjugated primary antibody that recognizes the 3X-FLAG tag on the N-terminus of the SMN transgenic construct. (B) Quantification of Western blot biological replicates represented in (A). Each sample contained 8-12 wandering 3^rd^-instar larvae, each genotype-temperature combination contains 3 to 5 samples (30-50 larvae total). Total protein was used as a loading control to standardize SMN levels (see Fig. S1). SMN levels for each sample were normalized to the “WT 25” SMN protein levels for their respective replicate. Error bars represent mean±95% c.i. Two-way ANOVA with Tukey post test yielded no significance.

To directly test the relative stabilities of WT and TDM SMN proteins in the absence of new protein synthesis, we carried out an *ex vivo* protein stability assay similar to the one described by Deliu et al., 2017. Wandering 3^rd^-instar WT and F70S animals (raised at 22°C) were dissected and then the larval filets were incubated in cell culture medium at 29°C in the presence or absence of cycloheximide (CHX) over a time course of 12 hours (Fig 4A). The F70S mutant was chosen for this assay because it displayed a pronounced temperature-sensitive phenotype. Samples were then analyzed for SMN protein levels via western blotting as shown in Fig. 4B. Puromycin was used as a secondary control to confirm that protein synthesis was effectively stalled in the presence of CHX (Deliu et al., 2017) (Fig S2). In untreated WT samples, SMN levels show a mild reduction from 0 to 12 hours post-exposure; however, in the presence of CHX, WT protein levels decrease consistently over the timecourse, illustrating the natural levels of SMN degradation during this time frame. In contrast, the F70S mutation causes SMN levels decrease more rapidly (Fig. 4B). Multiple replicates of each genotype at each timepoint verify these trends (Fig 4C), confirming that the F70S TDM is significantly less stable than its WT counterpart. These data provide the first evidence showing that SMN Tudor domain mutations trigger protein instability *in vivo*, providing molecular insight into mechanisms by which these point mutations can cause SMA. Additionally, the temperature-sensitive nature of these alleles provides a powerful tool for temporally regulating SMN protein levels throughout development and lifespan.

**Fig. 4.**
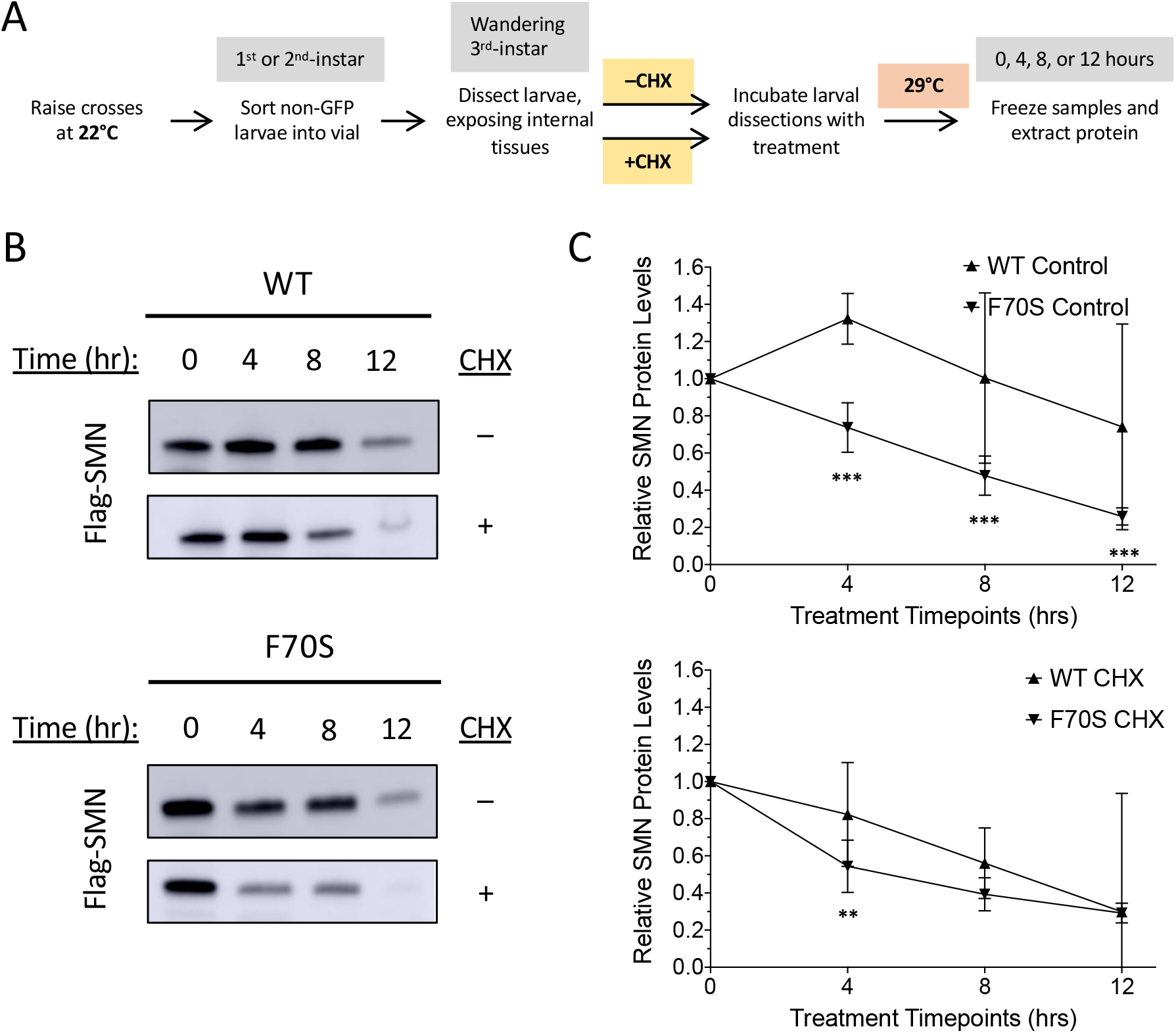
Cycloheximide experiments measure SMN protein stability in F70S Tudor domain mutants. (A) Schematic of cycloheximide (CHX) experiment, showing treatment types, timepoints, and treatment temperature (B) Representative Western blots of WT and F70S mutant wandering 3^rd^-instar larvae in the presence of CHX over 12 hours, 10 dissected larvae per sample. The control treatment contained Schneider’s media while the CHX treatment contained Schneider’s media and cycloheximide. All blots show FLAG-SMN levels, total protein used for standardization not shown. (C) Quantification of Westerns represented in (B), 3-9 samples per condition. “Control” represents the media only treatment, “CHX” represents the media and cycloheximide treatment. All SMN protein levels standardized using total protein, and relative to “0 hour” SMN levels of each genotype. Error bars represent mean±95% c.i. Adjusted P-values calculated using Tukey’s multiple comparisons test. ** p<0.01, *** p<0.001.

### High levels of SMN are not required during normal larval development but are essential for metamorphosis into adults

Numerous studies in SMA patients and animal models have shown that high SMN levels are vital during the early stages of development (Govoni et al., 2018; Jablonka and Sendtner, 2017; Ramos et al., 2019). Consistent with this idea, we previously showed that null mutations in *Drosophila Smn* result in developmental arrest and early larval lethality (Garcia et al., 2013; Praveen et al., 2012; Rajendra et al., 2007). Maternally deposited SMN is exhausted shortly after the first larval instar (Praveen et al., 2012), but a detailed analysis of SMN levels during later stages of development has not been performed. We therefore mined transcriptomic, proteomic and chromatin packaging databases to analyze expression from the *Smn* locus over developmental time. Furthermore, we exploited the temperature sensitivity of TDMs to help determine the requirements for high levels of SMN during larval, pupal and adult stages of *Drosophila* development.

As shown in Fig. 5A, developmental proteomic analysis shows that SMN protein is nearly 300-fold greater in embryos than it is during larval stages (Casas-Vila et al., 2017). Remarkably, chromatin accessibility (FAIRE-seq) analysis (McKay and Lieb, 2013) reveals that the *Smn* promoter region is essentially closed throughout, embryonic development, suggesting its transcriptional quiescence (Fig. 5B). Indeed, transcriptomic (RNA-seq) profiling of the same embryos shows that *Smn* mRNA levels progressively decrease during embryogenesis (Fig. 5B,C)(Graveley et al., 2011). We conclude that nearly all of the *Smn* mRNA and protein present in the animal during its first 24 hr of life is likely maternally deposited. Western blot analysis of *Smn* null mutants during early larval development showed that the maternal contribution of SMN protein persists throughout the first larval instar (L1), and is essentially depleted by the second, L2 (Praveen et al., 2012). These data are completely consistent with the developmental proteomics (Fig. 5A), showing that SMN levels during L2, early L3 and late (wandering) L3 are at or below the limits of detection (Casas-Vila et al., 2017). Moreover, SMN levels in newly-eclosed females are also very low but rise dramatically in 1-week old adults (Fig. 5A), again suggesting that the high levels of SMN detected in the older females is due to the production of eggs.

**Fig. 5.**
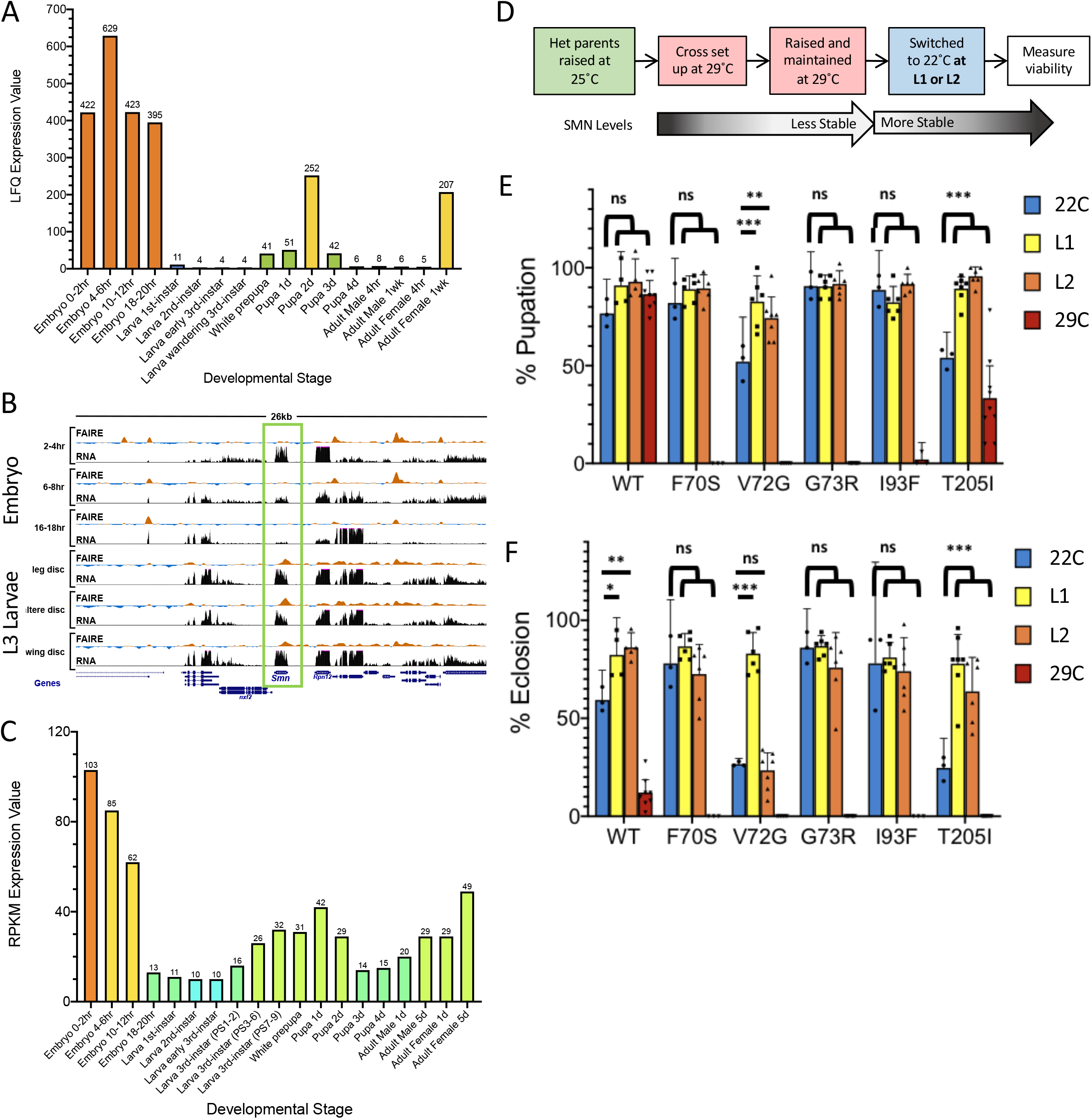
Nonpermissive-to-permissive temperature shift viability assays during early larval development. (A) Proteomic analysis of SMN levels throughout *Drosophila* developmental stages, data mined from Casas-Vila et al., 2017 via FlyBase (https://flybase.org/) (B) FAIRE and RNAseq analysis of *Drosophila Smn* genomic region over embryonic development and in larval imaginal discs, data mined from McKay and Lieb, 2013 (C) Transcriptomic analysis of *SMN* mRNA levels throughout *Drosophila* developmental stages, data mined from Graveley et al., 2011 via FlyBase (https://flybase.org/) (D) Workflow schematic of nonpermissive-to-permissive temperature shift experiments, switching larvae from 29°C to 22°C at 1 or 2 days post-egg laying (DPE). (E,F) Pupation (E) and eclosion (F) percentages of wildtype, Tudor domain mutant, and YG box mutant (T205I) larvae raised exclusively at 22°C (blue), switched from 22°C to 29°C at L1 stage (yellow), switched from 22°C to 29°C at L2 stage (orange), or raised exclusively at 29°C (red), 150-350 larvae per genotype, 50 larvae per biological replicate. Error bars represent mean±95% c.i. Adjusted P-values calculated using Tukey’s multiple comparisons test. ns: not significant (p >0.05), * p<0.05, ** p<0.01, *** p<0.001).

Transgenic lines expressing either *Smn* TDMs or controls were initially raised at 29°C (permissive), then switched to 22°C (non-permissive) at 1- or 2-days post-egg laying (DPE) and viability was assessed (Fig. 5D). Somewhat surprisingly, the large maternal contribution of SMN appears to be sufficient for embryonic development, even at the non-permissive temperature (Figs. 5E, F). Note, because SMN is required for ovarian development (Lee et al., 2009), we are unable to generate animals that completely lack maternal SMN. However, females that are raised at permissive temperature and then switched to the non-permissive temperature for mating and egg-laying are able to produce viable offspring if the progeny are switched to permissive temperature at either L1 or L2 (Fig. 5F). Control TDM larvae that are maintained at 29°C for the entire timecourse failed to pupate (Fig. 5E). Thus, as suggested by the proteomic data (Fig. 5A), high levels of SMN do not appear to be required for progression from L1 to L3.

Interestingly, the expression of SMN rises dramatically during pupation, only to drop again during later stages (Fig. 5A). In preparation for this second burst of activity during metamorphosis, the *Smn* promoter region is largely nucleosome-free by the time animals reach the wandering third (L3) instar and remains open in the pharate adult (Fig. 5B). To determine whether high levels of SMN are required for larval progression, pupariation and eclosion, we carried out temperature switch experiments. Progeny were initially raised at 22°C and then switched to 29°C after they reached the wandering 3^rd^-instar (Fig 6A). TDM larvae exposed to the non-permissive temperature at this later stage of development experienced significantly reduced pupation (Fig 6B). Even more striking, eclosion frequencies of the TDM larvae were similar to those of larvae that had been raised exclusively at 29°C (Fig 6C). These results demonstrate that although elevated SMN levels are not required for the earliest stages of larval development, high levels are required in order to complete metamorphosis.

**Fig. 6.**
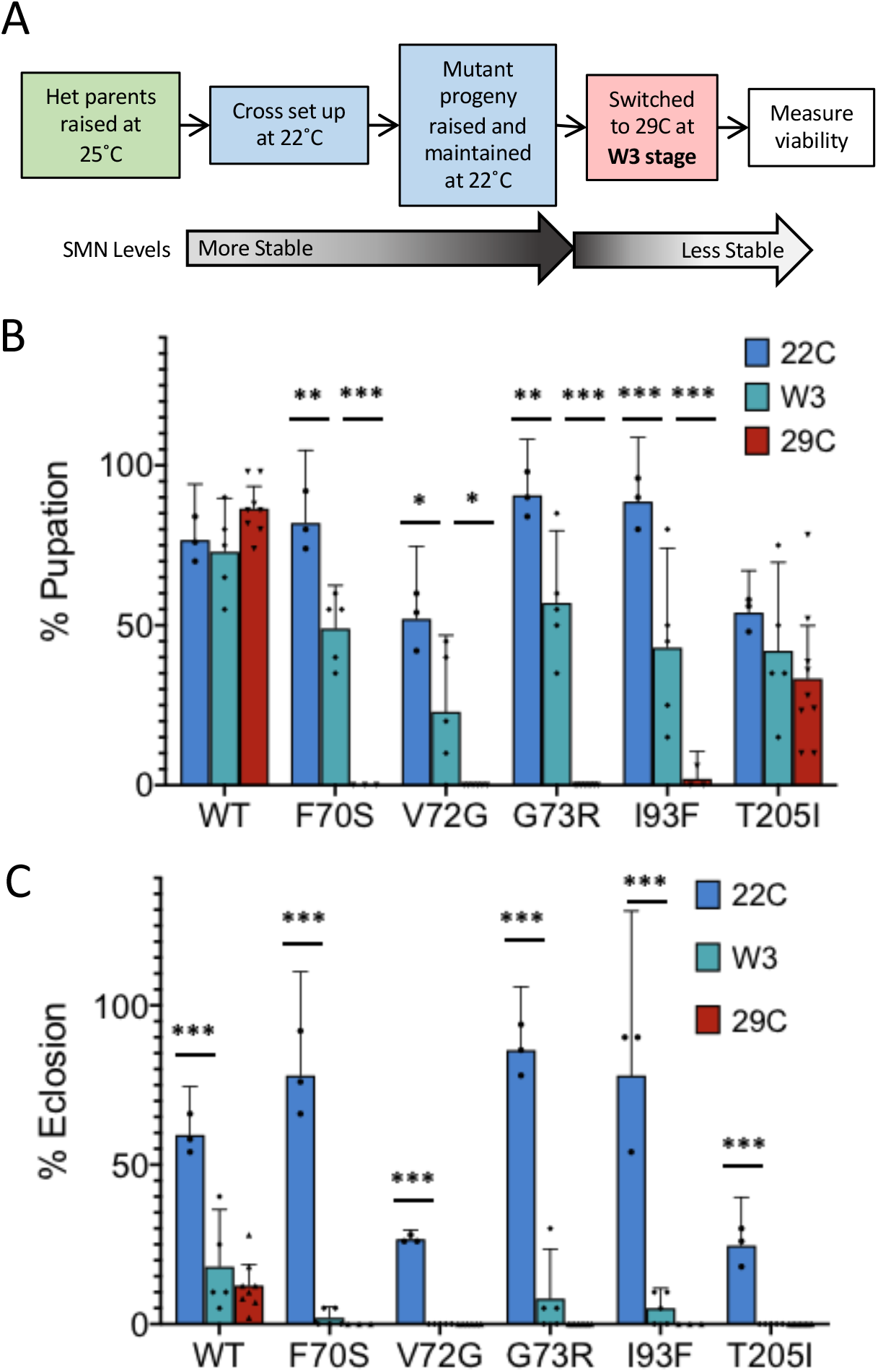
Permissive-to-nonpermissive temperature shift viability assays during late larval and pupal development. (A) Workflow schematic of permissive-to-nonpermissive temperature shift experiments, switching wandering 3^rd^-instar (W3) larvae from 22°C to 29°C. (B,C) Pupation (B) and eclosion (C) percentages of wildtype, Tudor domain mutant, and YG box mutant (T205I) larvae raised exclusively at 22°C (blue), switched from 22°C to 29°C at W3 stage (teal), or raised exclusively at 29°C (red), 150 larvae per genotype, 50 larvae per biological replicate. Error bars represent mean±95% c.i. Adjusted P-values calculated using two-way ANOVA and Tukey’s multiple comparisons test * p<0.05, ** p<0.01, *** p<0.001.

### Baseline levels of SMN are required for normal adult longevity

Finally, we examined the effect of decreased SMN stability during adulthood. Studies in mice show that high SMN levels are not required for survival as adults, however a baseline level is necessary for normal longevity (Sahashi et al., 2013). To test if the same trend holds in the fly, a subset of homozygous *Smn* transgenic stocks containing the WT, F70S, G73R, or I93F transgenes (as described in Materials and Methods) was used. These lines were raised through embryonic, larval, and pupal development at 25°C, with the exception of F70S, which had to be raised at 22°C in order to produce a testable number of adults. Within the first 24 hours post-eclosion, adult animals were either maintained at 25°C or switched to 29°C. This paradigm allowed us to assess the effects on longevity when SMN protein is reduced exclusively during the adult stage. Adult longevity was measured by recording the number of surviving adults every 2 days post-eclosion (Fig. 7A,B). As expected, all genotypes, including WT, showed reduced longevity at 29°C (Fig. 7A) compared to 25°C (Fig. 7B,C). However, the TDMs that were switched to non-permissive temperature experienced a significant drop in survival (dying within 6-16 days) compared to their permissive temperature counterparts (24-38 days) (Fig. 7A,B). To account for the baseline effects of elevated rearing temperature, we compared survival at 25°C and 29°C for each genotype (ratio: days to 10% survival at 29°/days to 10% survival at 25°C) (Fig. 7D). Importantly, when comparing the relative survival at the 10% survival threshold, the longevity of the WT line was 56% of that at 25°C. The G73R and I93F TDMs show a significant difference in longevity between these temperatures compared to WT, displaying 29°C longevities 32% and 35% the length of the respective longevities at 25°C (Fig. 7D).

**Fig. 7.**
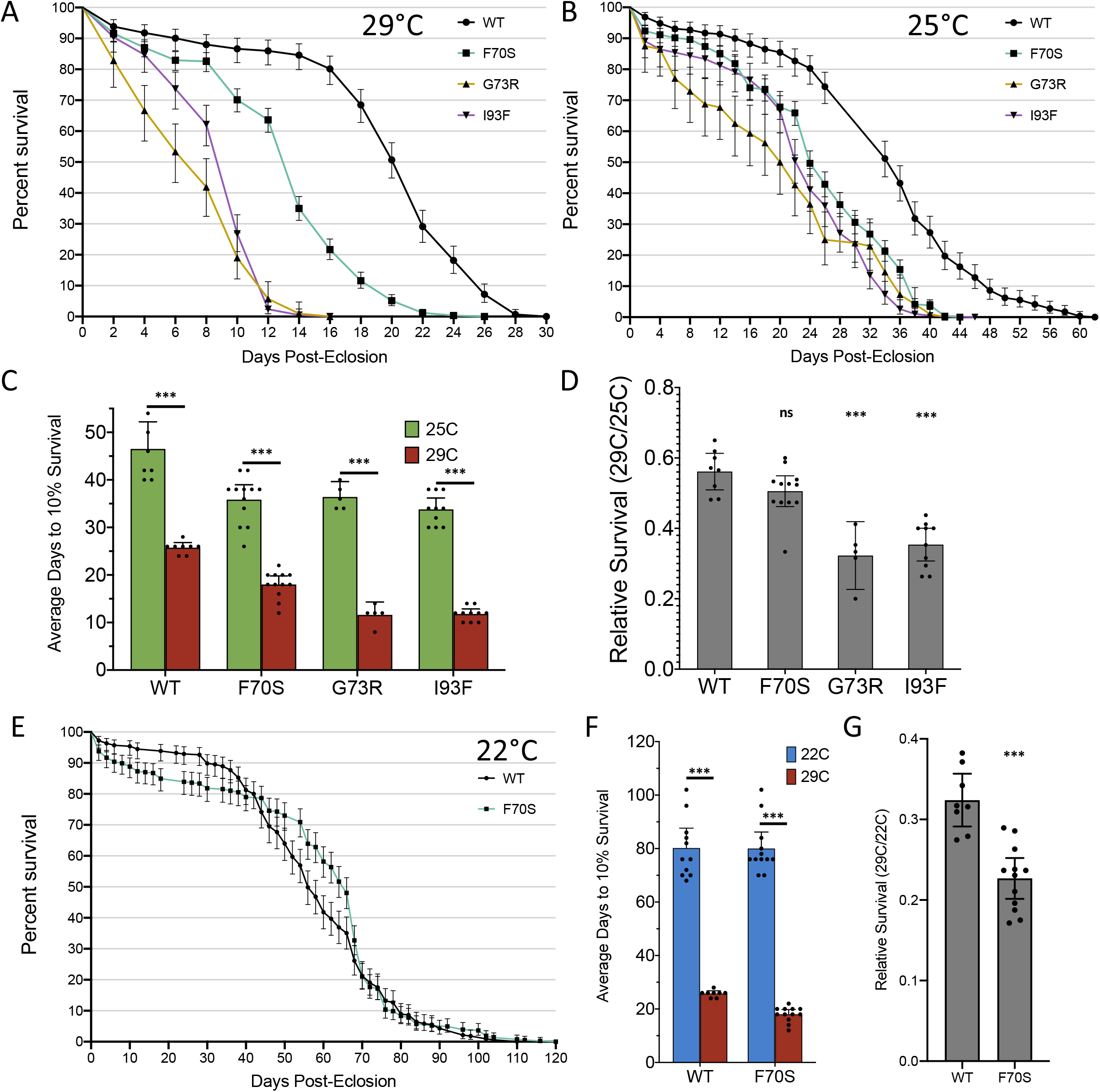
Adult longevity of select SMN Tudor domain mutants at permissive and non-permissive temperatures. (A,B) Survival plots of WT and Tudor domain mutant adult flies at (A) 29°C or (B) 25°C. The flies were raised at 25°C (WT, G73R, I93F) or 22°C (F70S) prior to eclosion and then moved to the experimental temperature <24 hours post-eclosion. The number of adults for each group range from 150 to 600. Live adults were counted every two days. Error bars represent mean±95% c.i. WT-black, F70S-green, G73R-purple, I93F-yellow. (C) Average time to reach 10% survival for adults at 29°C (red) and 25°C (green). Error bars represent mean±95% c.i. (D) Comparing relative longevity (10% survival 29°C/10% survival 25°C). Error bars represent mean±95% c.i. (E) Survival plot of WT (black) and F70S mutant (green) adult flies raised and maintained at 22°C (F) Average time to reach 10% survival for adults raised at 22°C (blue) and 29°C (red) Error bars represent mean±95% c.i. (G) relative survival (10% survival 29°C/10% survival 22°C). Error bars represent mean±95% c.i. P-values for 10% survival calculated using two-way ANOVA with Sidak’s multiple comparisons test. *** p<0.001. P-values for longevity ratios calculated using one-way ANOVA and student’s t-test. ns: not significant (p >0.05), *** p<0.001.

The F70S mutant showed 51% relative survival that was not significantly different from WT (Fig. 7D). This surprising result is hypothesized to be due to the fact that the F70S adults remain severely affected at 25°C (Fig. 1B,C), whereas the G73R and I93F defects at 25°C are mild. To account for this, a second study was carried out at 22°C with the WT and F70S transgenic stocks (Fig. 7E). Unlike other culturing temperatures, the WT and F70S longevity was similar at 22°C. When comparing the relative survival of these genotypes between 29°C and 22°C, there is a statistically significant difference in that the WT longevity displays milder changes in longevity (32%) than the F70S TDM (23%). F70S adults also showed a sex-specific difference in longevity that was not seen in the WT adults (Fig. S3). These data indicate that the TDMs have a differential sensitivity to temperature even after development is complete, and that an above-baseline level of SMN is needed into adulthood. Overall, these temperature-sensitive TDM lines can be utilized to further study the molecular, physiological, and behavioral consequences of modulating SMN protein levels in a living, developing organism.

## DISCUSSION

In this study, a series of point mutations within the SMN Tudor domain were found to exhibit pronounced temperature sensitivity relative to the wildtype or YG-box mutant SMN proteins. This differential sensitivity leads to significantly reduced SMN protein levels in the TDMs at higher temperatures. We utilized this paradigm to assess the effects of temporally manipulating SMN protein levels across various developmental timepoints.

### Effects of Tudor domain mutations on SMN stability

SMN protein levels are the strongest known modifiers of SMA disease severity, and small changes in these levels can dramatically affect age-of-onset and symptomatic severity. The relationship between SMN levels and SMA severity can be described using a two-threshold model (O’Hern et al., 2017). At levels above the upper threshold of SMN protein, individuals are unaffected. Below the second (lower) threshold, organisms die very early in development. Between these two thresholds of SMN expression is a region termed the “SMA zone,” wherein small changes in SMN levels cause large changes in disease severity (O’Hern et al., 2017).

Here, we show that single residue substitutions in the SMN Tudor domain directly impact SMN protein stability *in vivo*. This finding has important implications for the effects of similar mutations on SMA patients. In *Drosophila*, TDMs exhibit reduced SMN levels, leading to defects in organismal viability, locomotion and lifespan when animals are raised at higher temperatures (27°C and 29°C.). This idea is consistent with data from SMA patients suggesting that certain TDMs lead to reduced SMN protein levels in human cells (Takarada et al., 2017). Indeed, the phenotype of the *Drosophila* F70S mutant at the non-permissive temperature more accurately aligns with that of the corresponding human *SMN1* missense mutation, W92S, which causes Type 1 SMA (Kotani et al., 2007). Thus, the acute sensitivity of the TDMs to small shifts in temperature highlights a previously unrecognized variable in SMN biology.

Comparing our phenotypic data with the SMN Tudor domain structure sheds light on the relative importance of specific regions of the protein on its overall stability. Previous studies implicated the YG box in regulating SMN protein levels, however this is thought to occur via a completely distinct mechanism. That is, self-oligomerization of SMN leads to sequestration of a ubiquitin-dependent degron motif located within the SMN C-terminal region (Cho and Dreyfuss, 2010; Gray et al., 2018). The only Tudor domain mutation we assayed that does not display differential temperature sensitivity is Y107C. Unlike the other TDMs we tested, Y107C is located within the dimethylarginine-binding “pocket” of the Tudor domain (Tripsianes et al., 2011), and likely impacts SMN’s ability to bind Sm proteins and other potential targets.

One outstanding question is how biochemical/biophysical properties of these TDMs affect protein stability. It is likely that some or all of these mutations cause misfolding of the normal tertiary structures within the Tudor domain (Fig 8A). Previous *in vitro* and *in silico* studies showed that certain TDMs lead to misfolding and decrease stability (Hossain and Hosen, 2019; Li, 2017; Sneha et al., 2018; Tripsianes et al., 2011). Our own modeling shows that most SMN TDMs cluster around the same structural motif and cause a steric clash within this region when mutated (Fig 8B). It is also possible protein instability is due to loss of stabilizing interactions with binding partners, or a combination of both. However, our findings represent the first *in vivo* studies to measure the relative stability of SMN TDMs. Moreover, our experimental system allows us to test the function of an individual SMN mutant in the absence of wildtype SMN. These findings could prove important when developing treatments aimed at increasing SMN levels in patients. For example, small molecules targeting the Tudor domain could potentially improve SMN stability and increase protein levels at the post-translational level.

**Figure 8:**
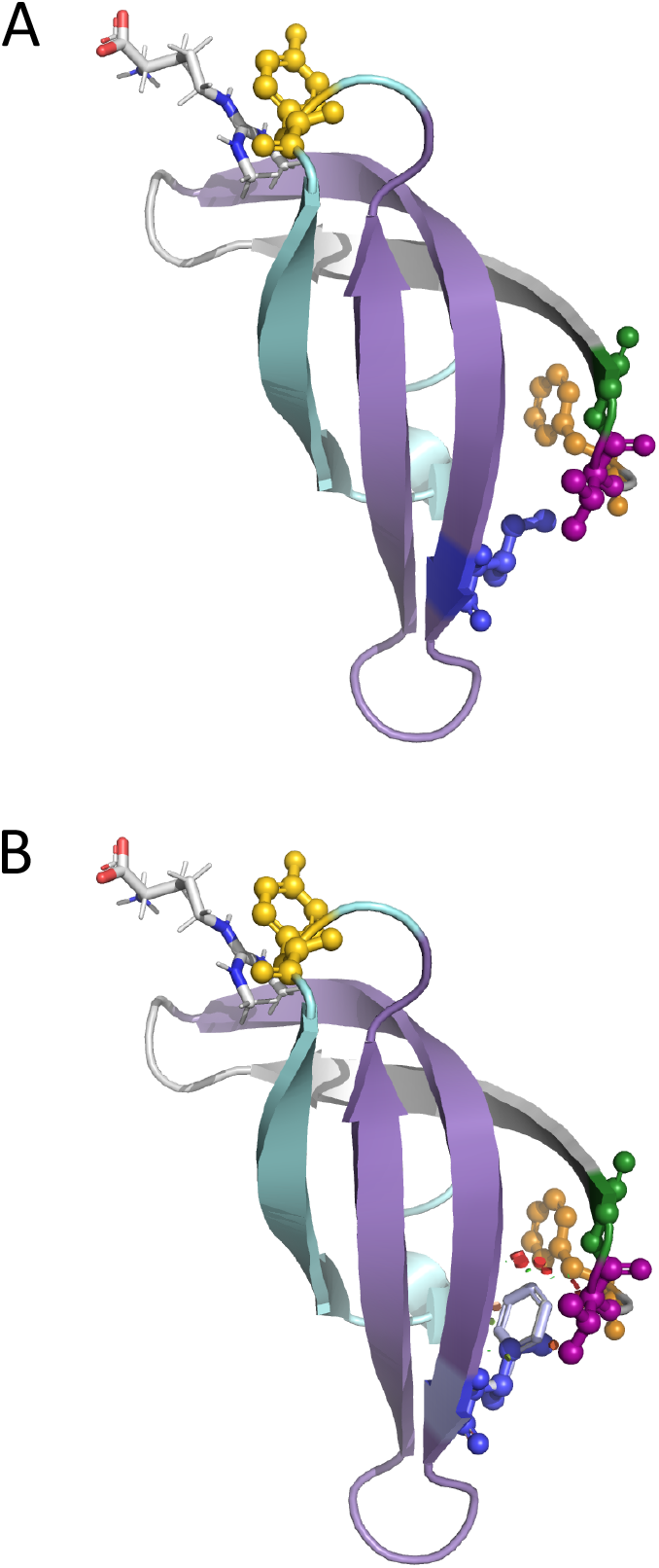
Structural consequence of SMN Tudor domain mutations. (A,B) Protein structure model of *Drosophila* SMN Tudor domain with bound dimethylated arginine (red-white-blue stick, upper left) (generation of model in Materials and Methods). (A) Wildtype SMN Tudor domain with residues of interest highlighted by ball-and-stick structures (orange-F70, magenta-V72, green-G73, blue-I93). (B) Same structure as (A) but with I93F overlay (silver). Steric clash shown by red circles. This is the most common of four conformations for I93F, but all four conformations experience steric clash.

### Temporal requirement for high SMN levels across development

In addition to uncovering a second disease mechanism, the discovery of temperature-sensitive SMN alleles provides a new genetic tool for the *in vivo* study of SMA-related phenotypes. Here, we have used this temporally-manipulatable system to test the requirements for high levels of SMN during *Drosophila* development. By manipulating the timing of exposure to permissive and non-permissive temperatures, we observed that production of high SMN levels is not required for larval development (1-2 DPE). Note that a baseline level of SMN is still required during these stages, as we and others have reported that *Smn* null animals undergo an early developmental arrest (Garcia et al., 2013; Shpargel et al., 2009).

We found that high SMN levels are crucial between the wandering 3^rd^-instar larval stage and the end of pupation. This finding correlates with whole-organism proteomic data showing that SMN levels are extremely high during both embryonic and pupal stages of *Drosophila* development (Casas-Vila et al., 2017). Interestingly, both embryonic and pupal stages involves the production of a new free-living animal. That is, embryogenesis results in a larva and metamorphosis involves a near complete regrowth of the body into an adult animal. As such, *Drosophila* pupal development more closely resembles peri-natal development in mice and humans. Our results show that it may be more medically relevant to directly compare developmental stages in which the majority of tissue formation and differentiation takes place. Thus, in terms of translational medicine, pupariation is perhaps a more appropriate stage of development on which to focus future studies of neuromuscular development.

### Temperature-sensitive SMN mutants as novel genetic tool for studying SMA

These temperature-sensitive SMN mutants can now be utilized as a system to further interrogate the effects of SMN protein level homeostasis at different stages of development. The ability to control SMN protein levels in our fruit fly lines using temperature allows us to more easily test molecular, physiological, and behavioral effects of reduced SMN levels during later developmental stages that were previously difficult to study because few individuals reached that stage. Similarly, we now have the ability to quickly rescue SMN protein levels simply by switching culturing temperatures. Altering SMN levels will enable us to study various longitudinal effects of SMN rescue at different points in development.

This model also provides a tool to screen for other factors involved in SMN biology and disease etiology. The TDM stable stocks can be successfully maintained at lower temperatures (22°C) and have the advantage of having no WT SMN, unlike the parental animals in a cross. These stable stocks are also beneficial because raising their culturing temperature can produce a situation where the majority of individuals die just before pupation or eclosion. When crossed to mutants of candidate genes or deficiency lines at these higher temperatures, any change in viability would be relatively easy to distinguish and signal a potential genetic interaction with *Smn*.

In conclusion, we have discovered a domain-specific effect on SMN stability that impacts SMN protein levels, motor function, and viability in *Drosophila* models of SMA. Further study of the Tudor domain and its role in the stability of SMN protein may be an important avenue for developing future SMA treatments that are effective in combination with existing therapies. The development of this temperature-sensitive model of SMA in *Drosophila* has allowed us to uncover the critical development timepoints for high SMN levels. In the future, this system could be used to further elucidate important functions of SMN during these stages and screen for novel disease interactors and pathways.

## MATERIALS AND METHODS

### Fly lines and husbandry

Balanced patient mutation lines (*Smn^X7^, Smn^TG^*/TM6B-GFP) were generated as described in Praveen et. al., 2012, where “TG” represents one of the fourteen *Smn* wildtype (WT) or mutant transgenes. Briefly, the lines were generated using ΦC31 integration at an insertion site located in chromosome band position 86F8. The *Smn* transgenic construct is a ∼3kb fragment containing the entire *Smn* coding region, expression of which is driven by the native *Smn* promoter. The transgene also contains an N-terminal 3X-FLAG tag that was used in this study to visualize SMN protein, circumventing the potential differences in α-SMN antibody binding between mutant forms of SMN. The *Smn^X7^* and *Smn^D^* alleles are previously described null alleles (Chang et al., 2008; Rajendra et al., 2007), and both stable stocks are GFP-balanced. To generate single-copy transgenic mutants (*Smn^X7^, Smn^TG^/Smn^X7^*) for the viability, locomotion, Western blot, and developmental timing assays, *Smn^X7^*/TM6B-GFP virgin females were crossed to *Smn^X7^, Smn^TG^*/TM6B-GFP males at the desired temperature (18, 22, 25, 27, or 29°C). Crosses were performed on molasses-based agar plates with yeast paste, and then GFP-negative larvae were sorted into vials containing standard molasses fly food at the 2^nd^-instar larval stage to prevent competition from heterozygous siblings.

The stable wildtype (WT) and mutant (F70S, G73R, and I93F) lines used in the longevity assays were generated by crossing *Smn^D^*/TM6B-GFP virgin females to *Smn^X7^, Smn^TG^*/TM6B-GFP males at room temperature. The progeny of these lines lacking the balancer chromosome were the allowed to propagate and develop into stable lines. The *Smn* copy number of individuals in these stocks is variable, with either one or both of the animal’s third chromosomes containing the transgene.

The WT and F70S lines used in the cycloheximide experiments were stable stocks homozygous for the transgene. These stock were generated by crossing males from the variable copy number stocks above to *Smn^X7^, Smn^TG^*/TM6B-GFP virgin females, and then sorting against the balancer chromosome markers.

All of the stocks except for the stable patient mutation lines were raised and maintained in a 25°C incubator unless being used for an assay. The stable patient mutation lines were raised and maintained at room temperature unless being used for an assay. Experimental temperatures for the assays were maintained using 18, 22, 25, 27, and 29°C incubators. All stocks were maintained in bottles containing standard molasses fly food.

### Viability assays

To assess viability, crosses were maintained and progeny were raised at the desired temperature on molasses agar plates. 25-50 GFP-negative progeny at the late 2^nd^ to early 3^rd^-instar stages were sorted into vials containing standard molasses fly food. After sufficient time had passed, pupal cases were counted and marked, and any adults were counted and removed from the vial. Any new pupal cases or adults were recorded every two days. The % viability was calculated at both the pupal and adult stages. Pupal viability (% pupation) was calculated by dividing the number of pupal cases by the initial number of larvae and multiplying by 100 (# pupae/# initial larvae*100). Adult viability (% eclosion) was calculated similarly but using the number of adults as the numerator (# adults/# initial larvae*100). Each vial was considered a biological replicate in respect to calculating averages and standard error.

### Larval locomotion assays

To assess the motor function of larvae at permissive and non-permissive temperatures, crosses were maintained and progeny were raised at desired temperature. Once the larvae reached wandering 3^rd^-instar larval stage, 1-5 larvae were placed onto the locomotion stage (a large molasses plate) at room temperature. The stage was then placed into a recording chamber to control light and reflections on the stage. Once all larvae were actively crawling, movement was recorded for at least 62 seconds on an iPhone6 at minimum zoom. Two recordings were taken for each set of larvae. At least 30 larvae were recorded for each experimental group. Locomotion videos were transferred to a PC and converted to raw video .avi files using the ffmpeg program. Videos were then opened in Fiji/ImageJ (https://imagej.net/Fiji), trimmed to about 60sec of video, and converted into a series of binary images. The wrMTrck plugin for ImageJ (http://www.phage.dk/plugins/wrmtrck.html) was used to analyze the video and determine larval size, average speed of movement, and average speed normalized to larval size (body lengths per second or BLPS) (Brooks et al., 2016). Each larvae was treated as an individual when calculating average and standard error.

### SMN Western blot analysis

To measure SMN protein levels of at different temperatures, 8-12 wandering 3^rd^-instar larvae were collected per sample, snap frozen in a dry ice ethanol bath, and stored at −80°C. Each sample was considered a biological replicate. Larval samples were then homogenized in RIPA buffer and 10X protease inhibitor cocktail (Halt*™* Protease Inhibitor Cocktail (100X), ThermoFisher) with a micropestle and spun at 13,000rpm at 4°C to separate the soluble phase. The supernatant was transferred to a new microcentrifuge tube and spun again to separate any lipid phase. The protein samples were then quantified using Bradford assay (BioPhotometer, eppendorf) and Western samples were prepped with 50ug of protein and 1X SDS loading buffer then denatured in a 95°C heat block for 5min.

Western samples were loaded and run in Mini-PROTEAN TGX Stain-Free Gels (BIO-RAD) at 300V for 15min (Mini PROTEAN*^®^* Tetra Cell, BIO-RAD). The total protein marker was UV-activated for 105 seconds (Fisher Biotech 312nm Transilluminator) and then the gel was placed in the transfer cassette (XCell II *™* Blot Module, novex *^®^* life technologies *™*) and transferred on to low-fluorescence PVDF membrane (Immun-Blot*®* LF PVDF Membrane Roll, BIO-RAD). After the transfer, total protein on the membrane was imaged using UV exposure with an Amersham Imager 600 (GE). The membrane was then blocked in 5% milk (in TBST) for 1hr at room temperature with gentle shaking, then incubated with α-FLAG HRP-conjugated primary antibody (1:10,000 in TBST; Monoclonal ANTI-FLAG*^®^* M2-Peroxidase (HRP) antibody produced in mouse, SIGMA) overnight at 4°C. The next day the membrane was washed 3 times for 5min in TBST at room temperature, then incubated with detection reagent (Amersham ECL*™* Prime Western Blotting Detection Reagents, GE) for 5min. The chemiluminescence was detected and imaged using an Amersham Imager 600 (GE). The FLAG-SMN levels and total protein were quantified using the ImageQuant TL 8.1 (1D analysis) program. Any samples with low-quality total protein signal were excluded from the analysis. Averages and standard error were determined based on the biological replicates for each condition. Any outliers were determined using the Grubbs’ test (https://www.graphpad.com/quickcalcs/grubbs1/) and removed from the data set before analysis.

### Protein stability assays

To assess the stability of SMN protein, cycloheximide treatment was applied to wandering 3^rd^-instar larvae. Larvae were produced from crosses and contained a single copy of the *Smn* transgene. Crosses and progeny were maintained at 22°C, where embryos were laid onto molasses-based agar plates. Larvae of the desired genotype were sorted into molasses food vials. When the larvae reached the wandering 3^rd^-instar developmental stage, they were dissected open to expose all the internal and external tissues to treatment media. The simple dissection was done by using dissecting tweezers to peel back a strip of the larva’s exoskeleton. 5 larvae were dissected for each sample. The control treatment contained 5ug/uL puromycin (SIGMA-ALDRICH, puromycin dihydrochloride from *Streptomyces alboniger* (P7255), stock solution of 25mg/mL dissolved in autoclaved water) in Schneider’s *Dorosophila* Medium (1X) (gibco, 21720-024). The experimental treatment contained 5ug/uL puromycin and 100ug/mL cycloheximide (SIGMA, C7698, stock solution of 10mg/mL dissolved in autoclaved water) in Schneider’s media. Larvae were placed in a microcentrifuge tube containing 500uL of treatment, and then nutated with the treatment for the desired time (hrs) at 29°C. After the desired treatment time, media was removed and sample preparation, protein extraction, and Western blot analysis were performed as described above. The α-puromycin antibody was used at a concentration of 1:1000 (Kerafast, #EQ0001), followed by an α-mouse secondary antibody at a concentration of 1:5000 (Pierce, PI31430).

### Temperature switch assays

To assess viability after exposure to permissive and non-permissive temperatures, SMN mutant larvae were switched between 22°C and 29°C at different developmental stages. The temperature switch assays in Figures 5 and 6 were performed with crosses, where mutant progeny contained one copy of the *Smn* transgene and a maternal component of wildtype SMN.

During the nonpermissive-to-permissive viability assays, crosses and progeny were maintained and raised at 29°C. Embryos were laid on molasses-based agar plates within a 4-hour time window, then either 1 or 2 days post-egg laying (DPE) the molasses plates were moved from 29°C to 22°C. Within 36 hours of being switched to 22°C, mutant larvae of the desired genotype were separated from their siblings and placed in a vial of molasses food (∼50 larvae/vial). Viability was then assessed by counting the number of pupae (% pupation) and adults (% eclosion) in each vial compared to the number of larvae as described above. Each vial was treated as a separate replicate and was used to calculate averages and standard error.

During the permissive-to-nonpermissive viability assays, the parental generation and progeny were raised at 22°C with the same molasses plate system and timed temperature switches as described above, with the single difference being that developing larvae were moved from 22°C to 29°C. In the wandering-3^rd^ temperature switch, progeny remained at 22°C and larvae were sorted into a molasses food vial while still at 22°C. Once the larvae began to wander, each larva was transferred to a new vial at 29°C (∼50 larvae/vial). Viability was assessed with the same method as above.

### Longevity assays

To assess longevity at permissive and non-permissive temperatures, newly eclosed adults (less than 24hr post-eclosion) were collected and put into fresh vials of molasses food. Males and females were separated into different vials. Each vial contained 10 or fewer adults to reduce stress and crowding. Around half of these adults remained at the permissive temperature (25°C or 22°) and the other half were switched to a non-permissive temperature (29°C). Animals were transferred to a fresh vial 2-3 times per week to prevent death due to sub-optimal food conditions. The number of surviving adults was recorded on the collection day and then every two days until all adults had expired. Any adults that were injured/killed or escaped during the experiment were removed from the counts. Every 20-50 adults were considered a biological replicate when determining averages, standard error, and survival thresholds.

### Structural Modeling

A model of dmSMN was generated using HHpred2 (Zimmermann et al., 2018). The template used was the Tudor domain from human SMN1 (PDB ID 4qq6) (Liu et al.). Figures of the 4qq6 structure and the dmSMN model were rendered in PyMOL [PyMOL The PyMOL Molecular Graphics System, Version 2.0 Schrödinger, LLC.]. A dimethylated arginine was placed in the active site of the 4qq6 and dmSMN structures based on the solution structure of the complex of human SMN with the dimethylated arginine ligand (PDB ID 4a4e) [PUBMED: 22101937]

### Statistical Analysis

All graphing and statistical calculations were performed using Prism (Version 8.2.0). All organismal viability, SMN protein levels (variable temperature and CHX), and temperature switch experiments were analyzed using a two-way ANOVA with Tukey’s multiple comparison test (alpha = 0.05). Larval locomotion was analyzed using unpaired multiple t-tests (alpha = 0.05). Adult longevity at 10^th^ percentile was analyzed using a two-way ANOVA with Sidak’s multiple comparisons test (alpha = 0.05). The 25°C vs. 29°C relative survival was analyzed using ordinary one-way ANOVA with Dunnett’s multiple comparisons test (alpha = 0.05). The 22°C vs. 29°C relative survival was analyzed using Welch’s t-test (alpha = 0.05). In all graphs, error bars are expressed as the mean±95% c.i.

## Acknowledgements

The authors are grateful to Dr. Brenda Temple and the R.L. Juliano Structural Bioinformatics Core facility at UNC-Chapel Hill for their expertise in protein structural modeling. We would also like to thank Dr. Joe Pearson for help with the developmental chromatin accessibility and gene expression profiling of the *Smn* locus. Finally, we would like to thank Drs. Harmony Salzler and Casey Schmidt for detailed editing of the manuscript.

## Author contributions

Conceptualization: ACR, ASM, AGM

Methodology: ACR, ASM, AGM

Formal analysis: ACR

Investigation: ACR, SSS, MRE, TPD, VV, ASM

Resources: AGM

Writing – original draft: ACR

Writing – review & editing: ACR, ASM, AGM

Visualization: ACR, AGM

Supervision: ACR, AGM

Project administration: AGM

Funding acquisition: AGM

---

**Supplemental Figure 1:**
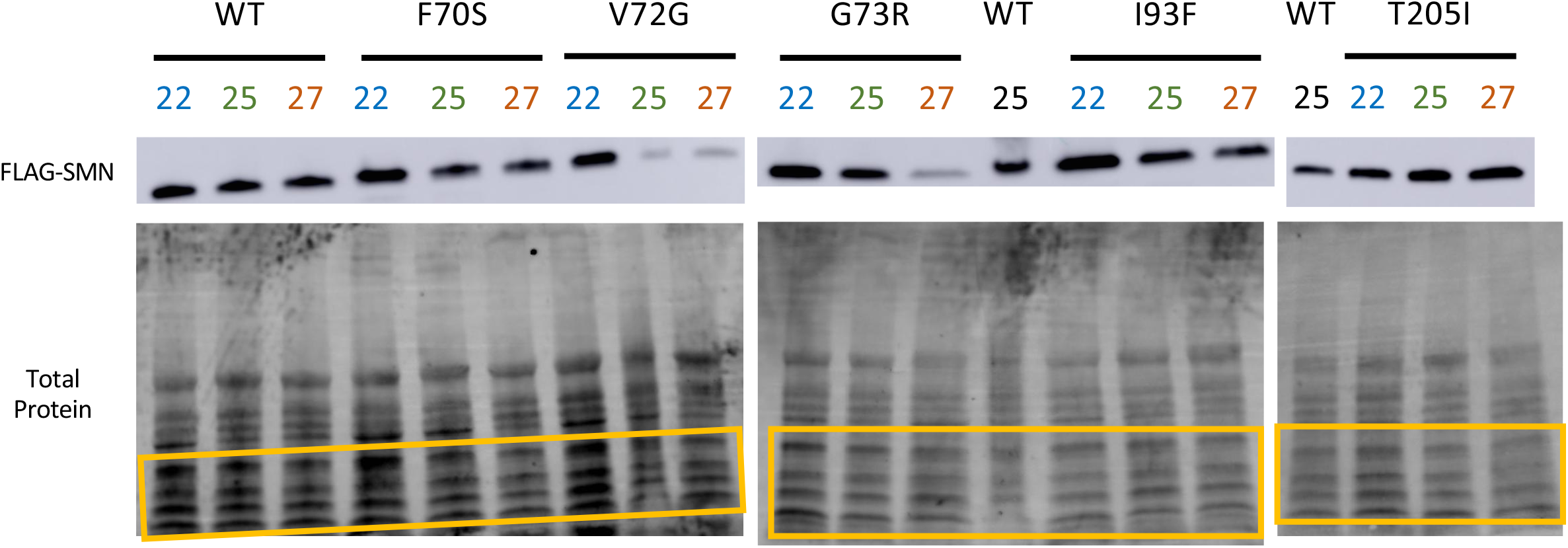
Representative FLAG-SMN Western blot and corresponding total protein UV fluorescence image. The FLAG-SMN blots are the same as in Figure 3. The total protein blot below each SMN blot is the same membrane detecting the total protein using UV fluorescence exposure. The yellow boxes distinguish the subset of four bands that was used in the quantification/normalization step of all Western blot analyses.

**Supplemental Figure 2:**
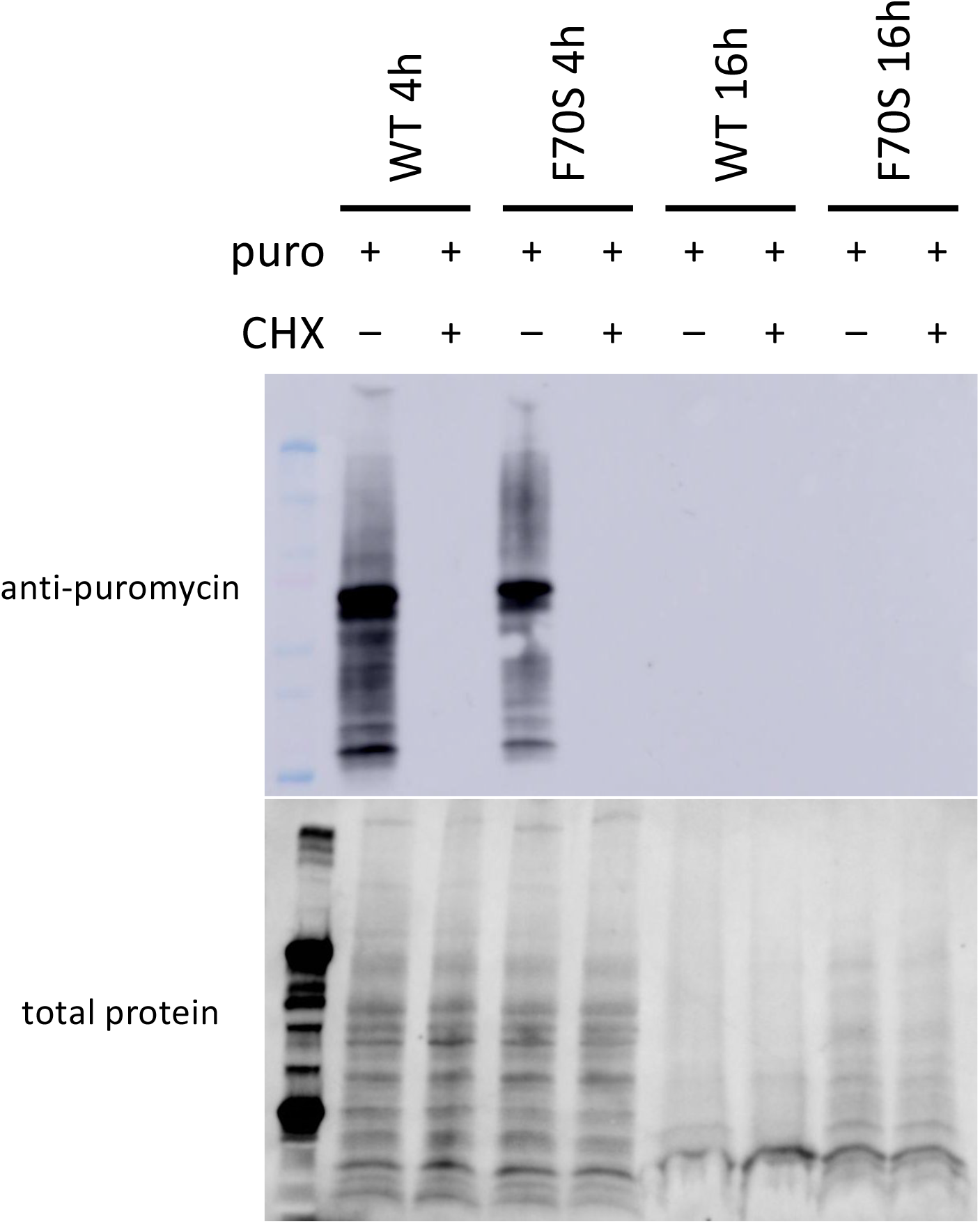
α-puromycin western blot in the presence and absence of cycloheximide (CHX). Western blot of wandering 3^rd^-instar wildtype larvae (10 larvae/sample) after puromycin treatment with or without CHX (4-hour or 16-hour treatment). Lower panel is total protein visualized using fluorescence as described in Materials and Methods. Upper panel is western blot of puromycin, which represents newly translated protein.

**Supplemental Figure 3:**
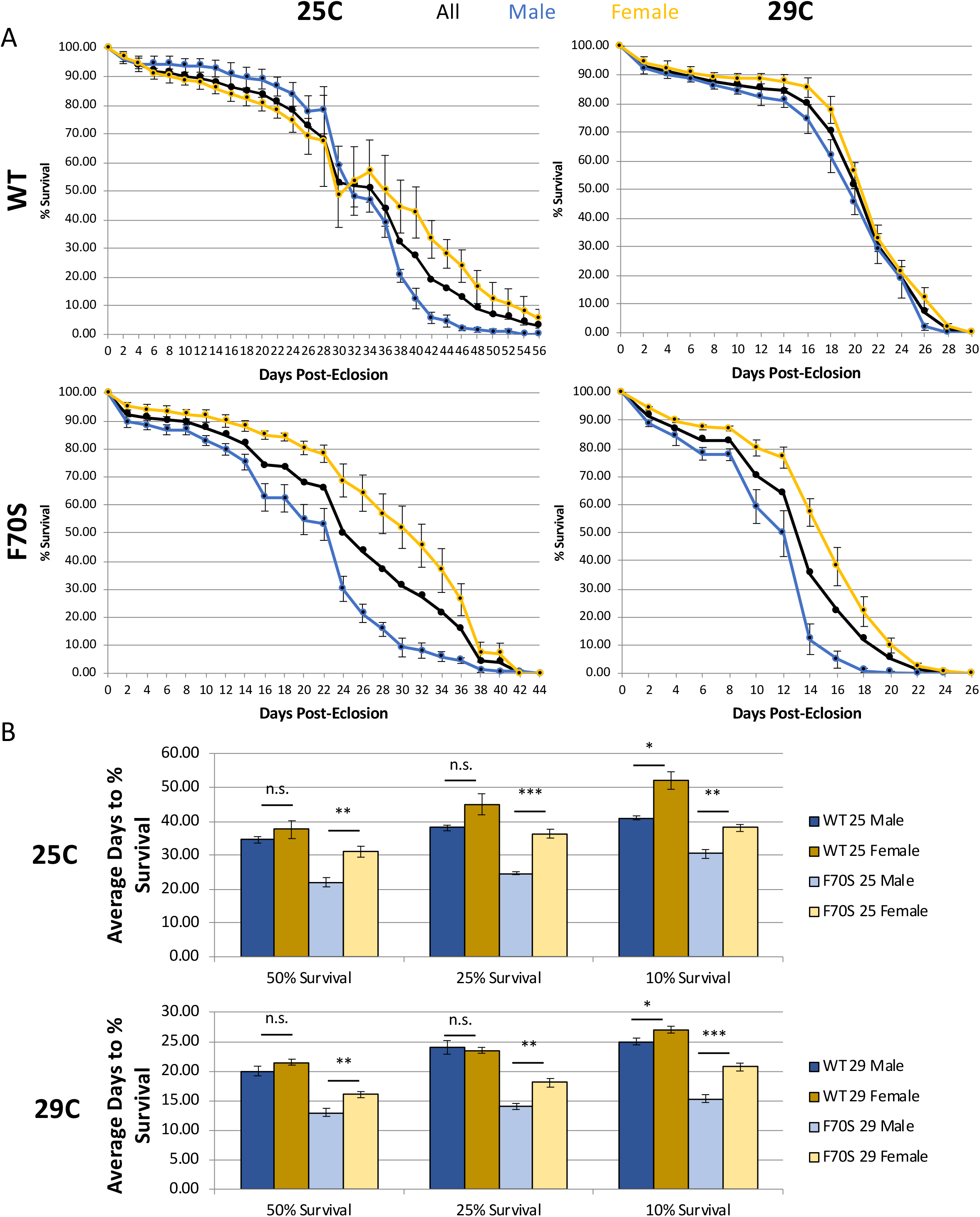
Sex-specific adult longevity. (A) Survival plots of WT and F70S Tudor domain mutant adult flies at 29°C (left) or 25°C (right). Number of adults for each genotype-temperature combination range from 140 to 300. WT survival plots are on top, F70S survival plots are one bottom. Live adults were counted every two days. Error bars represent standard error. Combined (Male+Female)-black, Male-blue, Female-yellow. (Left) WT 25°C flies were raised at 25°C and were shifted to a new vial remaining at 25°C until death; F70S flies were raised at 22°C and shifted to 25°C after eclosion. (Right) WT 29°C flies were raised at 25°C (F70S flies were raised at 22°C) and then both genotytpes were shifted to a new vial at 29°C less than 24h after eclosion until death (B) Comparing days to 50%, 25%, or 10% survival or less between male (blue) and female (yellow) adults at 25°C (above) or 29°C (below). Darker bars represent WT adults, lighter bars represent F70S adults. Error bars represent mean±s.e.m. P-values calculated using student’s t-test (* p<0.05, ** p<0.01, *** p<0.001).

